# Prenatal Stress Alters Transcription of NMDA-Type Glutamate Receptors in the Hippocampus

**DOI:** 10.1101/2024.02.18.580903

**Authors:** Tristram Buck, Erbo Dong, Alessandro Guidotti, Monsheel Sodhi

**Affiliations:** Department of Molecular Pharmacology and Neuroscience, Stritch School of Medicine, Loyola University Chicago, Maywood, IL. 60153.; Department of Psychiatry, Ohio State University, Columbus, OH.; Department of Psychiatry, University of Illinois at Chicago, Chicago, IL.

## Abstract

Prenatal stress damages the development of the cortico-hippocampal circuit in the brain and increases the risk for neurological disorders associated with deficits of social behavior, including schizophrenia. Accumulating evidence indicates that the NMDA-type glutamate receptor plays an important role in social cognition and stress-induced pathology in the hippocampus. In this study we have tested the hypothesis that transcription of NMDAR subunits is modified in the frontal cortex and hippocampus of prenatally stressed mice. Prenatal stress exposure was conducted by exposing pregnant mice to restraint stress three times daily during gestational weeks 2 and 3. We treated the adult offspring with haloperidol (1mg/kg), clozapine (5mg/kg) or vehicle (saline) twice daily for 5 days, after which we measured social interaction behavior (SI) and locomotor activity. After euthanasia, we measured the transcription of NMDAR subunits in the hippocampus and frontal cortex. We observed that saline-treated prenatally stressed (PRS-Sal) mice had reduced social interaction (SI) behavior compared to controls (NS-Sal) (P<0.01). This deficit was recovered in PRS mice treated with clozapine (PRS-Clz) but not the haloperidol-treated PRS group (PRS-Hal). These changes were not due to suppressed locomotion as neither PRS nor antipsychotic treatment reduced locomotor activity. These effects of prenatal stress were associated with increased transcription of NMDAR subunits (GRIN genes) in the hippocampus but not the frontal cortex. We observed positive correlation between GRIN transcription and social behavior in the frontal cortex, and conversely, negative correlation between GRIN transcription and social behavior in the hippocampus. Studies indicate that transcription of NMDARs is activity dependent, therefore altering the transcription levels of different NMDAR subunits would have a significant impact on the excitatory transmission in the corticolimbic circuit. The results suggest a molecular pathway by which prenatal stress in mice leads to life-long deficits in social behavior. It’s worth noting that while these associations have been observed in mice, the direct translation to human prenatal stress and NMDA receptor alterations requires further investigation. Nevertheless, these findings contribute to our understanding of the impact of prenatal stress on pathology in the hippocampus and downstream effects on social behavior and may have implications for understanding neuropsychiatric disorders related to prenatal stress exposure.

## Introduction

Prenatal stress impairs brain development of the fetus, and increases the risk for neurodevelopmental disorders (Khashan et al., 2008; Kinney et al., 2008; Li et al., 2010; Brown, 2011; Class et al., 2014; Veenstra-VanderWeele and Warren, 2015; Goldstein, 2019; Hicks et al., 2019). Prenatal restraint stress induces molecular pathways leading to structural deficits in the brain (Hayashi et al., 1998; Lemaire et al., 2000; Harrison, 2004; Charil et al., 2010; Miyagawa et al., 2011; Laloux et al., 2012; Gao and Penzes, 2015) but we have a limited understanding of the molecular mechanisms underlying these pathological effects(Son et al., 2006; Bustamante et al., 2010; Belnoue et al., 2013; Zhao et al., 2013; Benoit et al., 2015; Negron-Oyarzo et al., 2015). In experimental animals, prenatal stress is associated with pathology of the limbic circuitry, leading to abnormal behaviours, including increased fear and social memory deficits in males (Adrover et al., 2015) but not females (Schulz et al., 2014). We and others have previously shown that prenatal restraint stress results in deficits in social behavior in mouse offspring and that these deficits were prevented by treatment with the antipsychotic drug, clozapine. The social deficits observed were associated with altered molecular pathways in the hippocampus (Dong et al., 2015a; Dong et al., 2015b; Bristow et al., 2021).

Accumulating data indicate that the glutamate system plays an important role in social behavior. Treatment with the NMDA-type glutamate receptor (NMDAR) antagonist drug phencyclidine produces a deficit of social behavior that is ameliorated by clozapine (Qiao et al., 2001). The NMDAR is an ionotropic membrane-bound receptor that mediates fast excitatory transmission in the brain. The receptor is a tetramer, comprising varying combinations of GluN1, GluN2 and GluN3 protein subunits, encoded by the GRIN1, GRIN2 and GRIN3 genes respectively. Studies of human post-mortem brain reveal altered transcription of GRIN subunits in the frontal cortex and hippocampus in patients with stress-associated psychiatric disorders that include symptoms of social withdrawal, including schizophrenia (Meador-Woodruff and Healy, 2000; Harrison et al., 2003; Vrajova et al., 2010), but we do not understand the molecular basis of stress-induced abnormalities of social behavior. In this study, we have tested if prenatal stress altered the transcription of NMDAR subunits in the prefrontal cortex or hippocampus in mice, and if antipsychotic drug treatment modify the transcription of NMDAR subunits. Our results show that prenatally stressed mice have higher levels GRIN2A and GRIN2B subunit transcription in the hippocampus but not in the prefrontal cortex, and that treatment with clozapine appears to prevent these transcriptional modifications. We also report differential correlation of the transcription of NMDAR subunits with behavior in the frontal cortex and hippocampus. These data indicate that transcription of the NMDAR subunits may play a role in the pathophysiology of prenatal stress and its long-term impact on behavior.

## Materials and Methods

### Animals and PRS Procedure

All procedures were performed according to NIH guidelines for animal research (Guide for the Care & Use of Laboratory Animals, NRC, 1996) and approved by the Animal Care Committee of the University of Illinois at Chicago and Loyola University Chicago. Pregnant mice (Swiss albino ND4, Harlan, Indianapolis, IN) were individually housed with a 12-h light-dark cycle and access to food and water *ad libitum.* Dams assigned to the stress group were subjected to repeated sessions of restraint stress, as described previously, while dams assigned to the control group were left undisturbed throughout gestation. The stress protocol (PRS) consisted of restraining pregnant dams in a transparent tube (12 × 3 cm) under bright light for 45 minutes three sessions a day from day 7 of pregnancy until delivery. After weaning (postnatal day/ PND 21), male offspring were housed in groups of 4-5 based on condition. Each treatment group (controls, prenatal stress) contained mouse offspring from an average of 6 different litters. The experimental design is summarized in **Figure 1A**, and we summarize the mouse subgroups in **Supplementary Table 1**.

**Figure 1:**
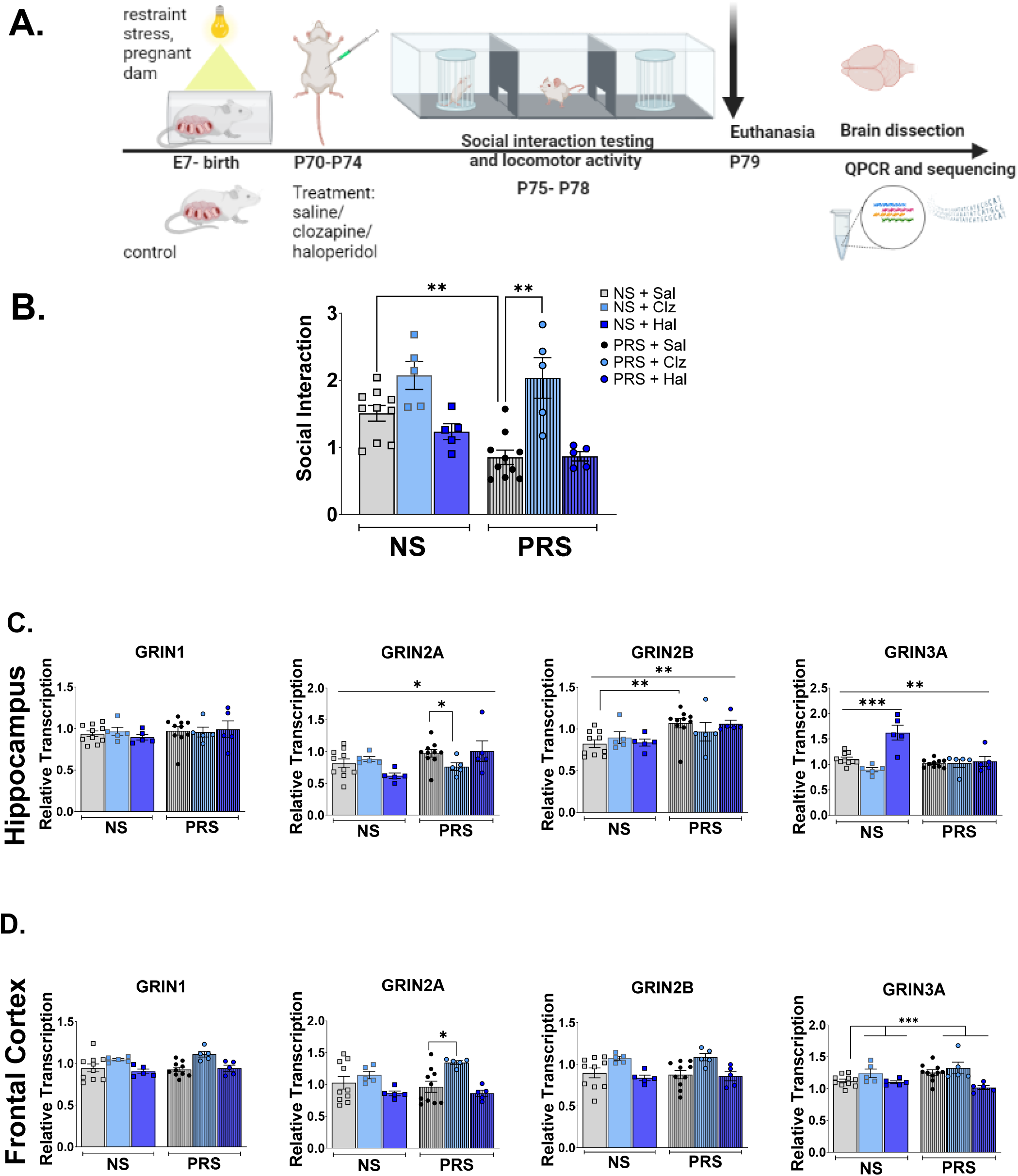
Prenatal Stress Induces Social Deficits and Alters GRIN2 Transcription in the Hippocampus and Frontal Cortex. **(A)** Schematic diagram of study protocol showing timeline of prenatal stress, drug treatment of offspring, analyses of offspring behavior and gene transcription. We injected mice offspring exposed to prenatal stress (PRS) and the non-stressed control group (NS) with vehicle, clozapine (5mg/kg), or haloperidol (1mg/kg) twice daily for 5 days from postnatal day 70. We subsequently tested locomotor activity and SI behavior. After euthanasia, we measured mRNA abundance of GRIN genes (see Methods). **(B)** The social interaction activity (ratio of sniffing time to the wire cup containing a stranger mouse vs sniffing time of the empty cup) was measured 18 hours after the last antipsychotic treatment. The data are expressed as mean ± sem. PRS causes a deficit of social interaction behavior that is not observed in PRS mice treated with clozapine. PRS mice had lower SI relative to NS mice (F_1,18_ =17.0, p=0.001). PRS mice treated with clozapine (PRS-Clz) had higher SI levels than the PRS group treated with vehicle (PRS-saline), (F_1,8_ =11.5, p=0.009), as previously reported (Dong et al., 2015a). In contrast, haloperidol did not alter SI in any group. **(C)** *Post-hoc* tests of the transcription of GRIN genes in the hippocampus showed that PRS mice had increased transcription of GRIN2A (F_1, 34_=4.30, p<0.05, η^2^=0.11), and GRIN2B (F_1, 34_=11.1, p=0.002, η^2^=0.25). In contrast PRS mice had decreased transcription of GRIN3A (F_1, 34_=9.89, p=0.003, η^2^=0.23) and no change in GRIN1 transcription relative to NS mice. Antipsychotic drugs modified GRIN3A transcription in NS but not PRS groups. Relative to saline-treated mice, clozapine treated mice had reduced GRIN3A transcription in the NS group whereas haloperidol treatment increased GRIN3A transcription (F_2, 34_=14.4, p=0.00003, η^2^=0.46). GRIN2A transcription was reduced in clozapine treated PRS mice relative to saline treated PRS mice (p<0.05). Antipsychotic drugs did not alter the transcription of other GRIN genes in the hippocampus. **(D)** *Post hoc* application of the non-parametric Welch’s test to the transcription of GRIN1, GRIN2A and GRIN2B genes revealed that only GRIN2A had differential transcription between subgroups of mice. We observed an increase of GRIN2A transcription in clozapine-treated mice relative to other groups (H_1, 5.81_=8.73, p=0.03). We were able to use ANOVA to analyze effects of stress and drug treatment on GRIN3A transcription in the frontal cortex, and we detected a significant effect of drug treatment but not stress on GRIN3A transcription (F_2, 34_=8.24, p=0.001, η^2^=0.33). This effect was due to reduced GRIN3A transcription in the haloperidol treated PRS mice. *Abbreviations:* Embryonic age E; Postnatal age P; Frontal Cortex, FCx; Glutamate Receptor Ionotropic NMDA, GRIN; non-stressed control, NS; prenatally stressed, PRS; clozapine treated, Clz; haloperidol treated, Hal; saline treated, Sal. *p≤0.05, **p<0.01, *** p<0.001.

**Table 1:**
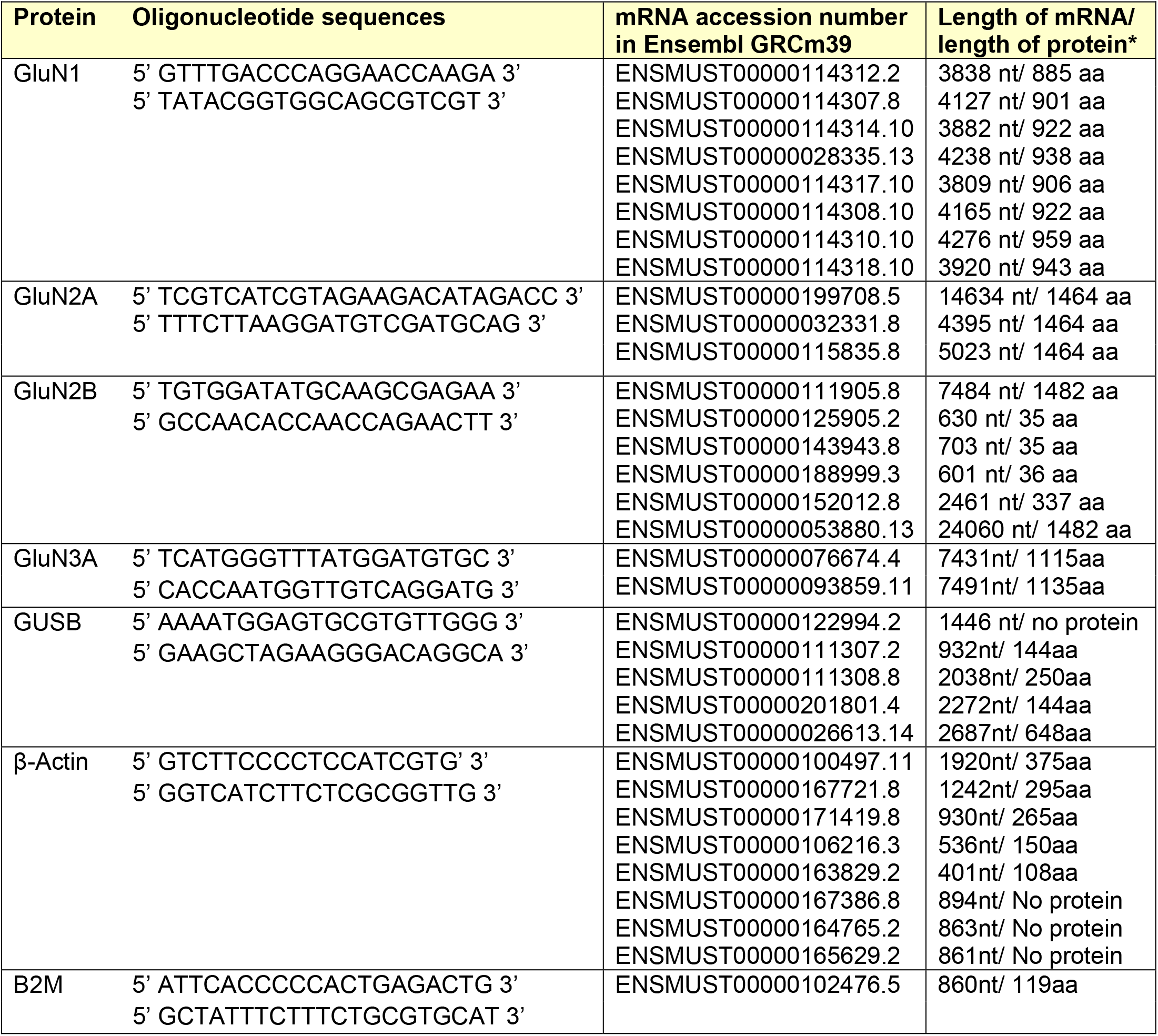
Assays designed to target mRNAs of NMDA receptor subunits that are translated. Specific mRNAs that are amplified by each primer pair are indicated. *nt, nucleotide; aa, amino acid.

### Drug Treatment

The drugs, haloperidol (Sigma, St Louis, MO) and clozapine (Novartis Pharmaceuticals, Basel, Switzerland) were prepared daily and dissolved in glacial acetic acid and sodium hydroxide (NaOH, Sigma) to reach pH 6. PRS and non-stressed (NS) control mice received subcutaneous injections of either haloperidol (1mg/kg), clozapine (5mg/kg) or saline (vehicle/ veh) twice daily from PND 70 for 5 days. Drug doses were determined based on previous research. The dose of haloperidol was selected according to dopamine D2 receptor occupancy, which is a predictor of antipsychotic efficacy for neuroleptic drugs (Wilmot and Szczepanik, 1989). The dose of clozapine tested has been shown to reverse the cognitive deficit induced by sub-chronic phencyclidine (Grayson et al., 2007) and aligns with the dosage range of clozapine used to treat schizophrenia (Subramanian et al., 2017). We began measuring mouse behavior 18 hours after the final injection. Behavioral tests included measures of locomotor activity, followed by social behavior tests, on consecutive days. We euthanized mice one day after the final behavioral test. We dissected and stored the brain tissue at -80 deg C.

### Locomotor activity

We tested locomotor activity and social interaction in succession on the same day at PND 75 between 10 am and 3 pm. We selected PND 75 for behavioral testing because offspring performance at PND 75 was more reproducible and stable than performance measured at earlier ages. Before the three-chamber test, we assessed if changes in SI might be confounded by locomotor activity. We used a computerized system with VersaMax software (AccuScan Instruments, Columbus, OH) to quantify and track locomotor activity in mice, as described previously (Carboni et al., 2004). A Perspex box (20×20×20cm divided into quadrants) was surrounded by horizontal and vertical infrared sensor beams. Activity was recorded for 15 minutes. The total number of interruptions of the horizontal sensors (counts) was taken as a measure of horizontal activity and counts of interruptions of vertical sensors were a measure of vertical activity (rearing).

### Social Interaction (SI) Behavior

We used the Three-Chambered Apparatus method to measure social interaction (SI) (Nadler et al., 2004; McFarlane et al., 2008; Dong et al., 2015a). The apparatus is a transparent acrylic, three-chamber box with each chamber measuring 20cm × 40.5cm × 22cm. The central chamber walls had openings that were 10cm × 5cm, allowing mice to move into the side chambers. The chambers on either side of the central chamber contained identical wire cups, but one of the cups contained a second mouse (a stranger of the same sex), while the other cup remained empty. In the initial phase of the assay, the test mouse explored the empty apparatus for 5 min. Subsequently, we confined the mouse to the central chamber and placed the stranger mouse in one of the wire cups, in a side chamber. The test began by allowing the test mouse to explore all three chambers freely for 10 min. Social interaction (SI) was defined as the ratio of the sniffing time at the cup containing the stranger mouse vs. the cup containing no mouse/ empty cup. Between tests the apparatus was washed with 70% ethanol and distilled water. We performed tests in dim lighting between 10am and 3pm, with sessions recorded for data analysis. Inter-rater reliability was assessed by correlating the scores of two raters observing the video recordings of the test (VersaMax, AccuScan Instruments, Columbus, OH).

### RNA Extraction and Gene Expression Analysis

We homogenized the whole hippocampus and extracted total RNA using TRIzol reagent (Invitrogen, Carlsbad, CA). We synthesized complementary DNA (cDNA) from 100 ng of RNA in a 40µl reaction containing 100 units of Tetro Reverse transcriptase (Bioline, Taunton, MA), 8µl of 5X RT buffer, 1mM dNTP Mix (Bioline), 40 units of Ribosafe RNAse Inhibitor (Bioline), and 4µl of 10X random primers (Applied Biosystems, Foster City, CA). The reaction cycling conditions were 25°C for 10 min, 37°C for 120 min, and 85°C for 10 min in a Veriti Thermal Cycler (Applied Biosystems).

We measured mRNA abundance of GRIN1, GRIN2A, GRIN2B and GRIN3A, normalized relative to the abundance of mRNAs encoding housekeeping genes β-Actin and β2 microglobulin (B2M) using quantitative polymerase chain reaction (qPCR). Each qPCR reaction included 0.8µl of cDNA and FastStart Universal SYBR Green Master (Roche, Basel, Switzerland) in a 12µl reaction. All primers were designed so that the sense and corresponding antisense primers were located in different exons (sequences listed in **Supplementary Table 2**), to prevent amplification of genomic DNA. QPCR included an initial denaturation step of 94°C for 5 min followed by 40 cycles of 94°C for 30 sec, 61°C for 30 or 45 sec, and 72°C for 30 sec. We performed assays in duplicate in 96-well optical plates using the MX3000P instrument (Stratagene, La Jolla, California) and Sequence Detector Software (SDS version 1.6; PE Applied Biosystems). We used the relative standard curve method for these analyses (https://assets.thermofisher.com/TFS-Assets/LSG/manuals/cms_042380.pdf).

### Statistical Analyses

We used SPSS version 24 (IBM, Armonk, NY) to perform statistical analyses. We tested for differences in gene transcription between treatment groups in each brain region using two-way multivariate ANOVA.

We tested if the transcription of genes encoding NMDAR subunits were altered by prenatal stress or antipsychotic drugs using the equation below:

Gene transcription (y) = β0 + β1*stress + β2*drug + β3*stress*drug

With stress as a two-level factor (prenatal stress, no stress), and drug as a three-level factor, (vehicle, haloperidol and clozapine).

We analyzed GRIN transcription for 40 mice exposed to one of two stress conditions: prenatal stress or no stress, and one of three drug treatments: saline, haloperidol or clozapine. Our aim was to discover whether GRIN expression in the hippocampus or frontal cortex differs based on mouse stress exposure and drug treatment. Our dependent variable was GRIN transcription, and our first independent variable was stress exposure, i.e. prenatal stress or no stress. Since stress has two levels and drug treatment has three levels, we had six groups of participants in our study. There were 10 mice in each stress group and 10 mice in each drug treatment group as indicated in **Supplementary Table 1**.

The two-way ANOVA tested three null hypotheses:

1. The main effect for the first independent variable (stress) on GRIN transcription.
*H_0_*: There is no difference between the mean GRIN transcription scores of PRS and NS mice.

2. The main effect for the second independent variable (drug treatment):
*H_0_*: There are no differences between the mean GRIN transcription for mice treated with different drugs.

3. Interaction effect:
*H_0_*: There is no interaction between stress exposure and drug treatment on mean GRIN transcription.

This third hypothesis tests whether the impact of drug treatment on GRIN transcription depends on the level of stress exposure i.e. is the effect of stress on mean GRIN transcription altered by antipsychotic drug treatment?

We tested each brain region separately (frontal cortex, hippocampus). In *posthoc* analyses, we tested the relationship between GRIN gene transcription and SI behavior using linear regression. Where data were not normally distributed, we used Spearman’s rank correlation analysis, Kruskal-Wallis, or Mann-Whitney U tests, as appropriate. False discovery rate (FDR) correction for multiple comparisons was calculated by the method of Benjamini and Hochberg (Benjamini and Hochberg, 1995).

## Results

As in previous studies (Dong et al., 2015a; Bristow et al., 2021), litters for Swiss Webster mice in our laboratory contain 8-10 pups. Body weights of offspring from PRS and non-stressed (NS) groups at postnatal day 75 (late adolescence–early adulthood in mice) were not significantly different (p>0.1). Vehicle-treated PRS mice had lower SI compared with vehicle-treated non-stressed mice in adulthood as previously reported by us (Dong et al., 2015a; Bristow et al., 2021).

### Prenatal stress reduced social interaction which was prevented by clozapine treatment

Data are summarized in **Figure 1B**. Levene’s test revealed that the error variance of social behavior differed between the stress and drug treatment groups (p<0.05). Kruskal-Wallis tests indicate that PRS reduced social interaction behavior in saline-treated (H_1_=20.7, p=0.000005) and haloperidol-treated mice (H_1_=8.31, p=0.004) but not in clozapine-treated mice. These behavioral effects of PRS are consistent with previous reports (Guidotti et al., 2014; Dong et al., 2016b; Matrisciano et al., 2018). PRS and drug treatment did not alter horizontal locomotor activity (p>0.05).

### Hippocampus: Transcription of GRIN genes altered by prenatal stress and antipsychotic drug treatment

#### Effect of prenatal stress (PRS) in the hippocampus

Two-way multivariate analysis of variance (MANOVA) for the effect of PRS on GRIN gene transcription (all genes) was significant, F_4, 31_=7.04, p<0.001, η^2^=0.48, Wilks’ Λ=0.524. Levene’s test of equality of error variances was not significant (p>0.05). We summarize *post-ho*c analyses of GRIN transcription data in **Figure 1C**.

#### Effect of antipsychotic drugs in the hippocampus

MANOVA showed a significant effect of antipsychotic drug treatment on GRIN gene transcription, F_8, 62_=4.271, p<0.001, η^2^=0.36, Wilks’ Λ=0.416. *Post-hoc* tests are summarized in **Figure 1C**.

#### Interaction of PRS and drug treatment in the hippocampus

MANOVA revealed a statistically significant interaction between drug treatment and PRS exposure on transcription of all GRIN genes measured in the hippocampus, F_8, 62_=4.461, p<0.001, η^2^=0.37, Wilks’ Λ=0.403. *Post-hoc* tests of individual GRIN genes showed an interaction of antipsychotic drug treatment and PRS on the transcription of GRIN2A (F_2, 34_=3.91, p=0.03, η^2^=0.19) and GRIN3A (F_2, 34_=11.5, p=0.0002, η^2^=0.40) but not GRIN1 or GRIN2B.

### Frontal cortex: GRIN transcription is altered by antipsychotic drug treatment and prenatal stress

Levene’s test revealed homogeneity of variances revealed that the error variance of GRIN1, GRIN2A and GRIN2B transcription across groups in the frontal cortex was not equal and therefore these values were not normally distributed. There was a medium effect of PRS on GRIN transcription in the frontal cortex (η^2^=0.08), and a large effect of antipsychotic drug treatment on GRIN transcription in the frontal cortex (η^2^=0.32). There was also a large effect on GRIN transcription in the frontal cortex due to the interaction of drug and stress exposure (η^2^=0.14). *Post-hoc* analyses of individual genes are summarized in **Figure 1D**.

### Relationship between GRIN transcription and social interaction behavior

We proceeded to test if there was a linear relationship between GRIN transcription and social behavior (see **Figure 2**). Normal probability plots of the residuals of social interaction with GRIN transcription in the hippocampus or frontal cortex of saline-treated mice (n=20) were approximately linear. This supported the condition that the error terms were normally distributed. Overall GRIN transcription in the frontal cortex did not have a linear relationship with social interaction behavior, and *post-hoc* analyses of individual GRIN genes showed a positive correlation with social interaction (**Figure 2A)**. GRIN transcription in the hippocampus had a linear relationship with social behavior (r=0.87, r^2^=0.76; F_4, 15_=11.8, p=0.00015). *Post-hoc* analyses of individual GRIN genes show a negative correlation with social interaction (**Figure 2B**).

**Figure 2:**
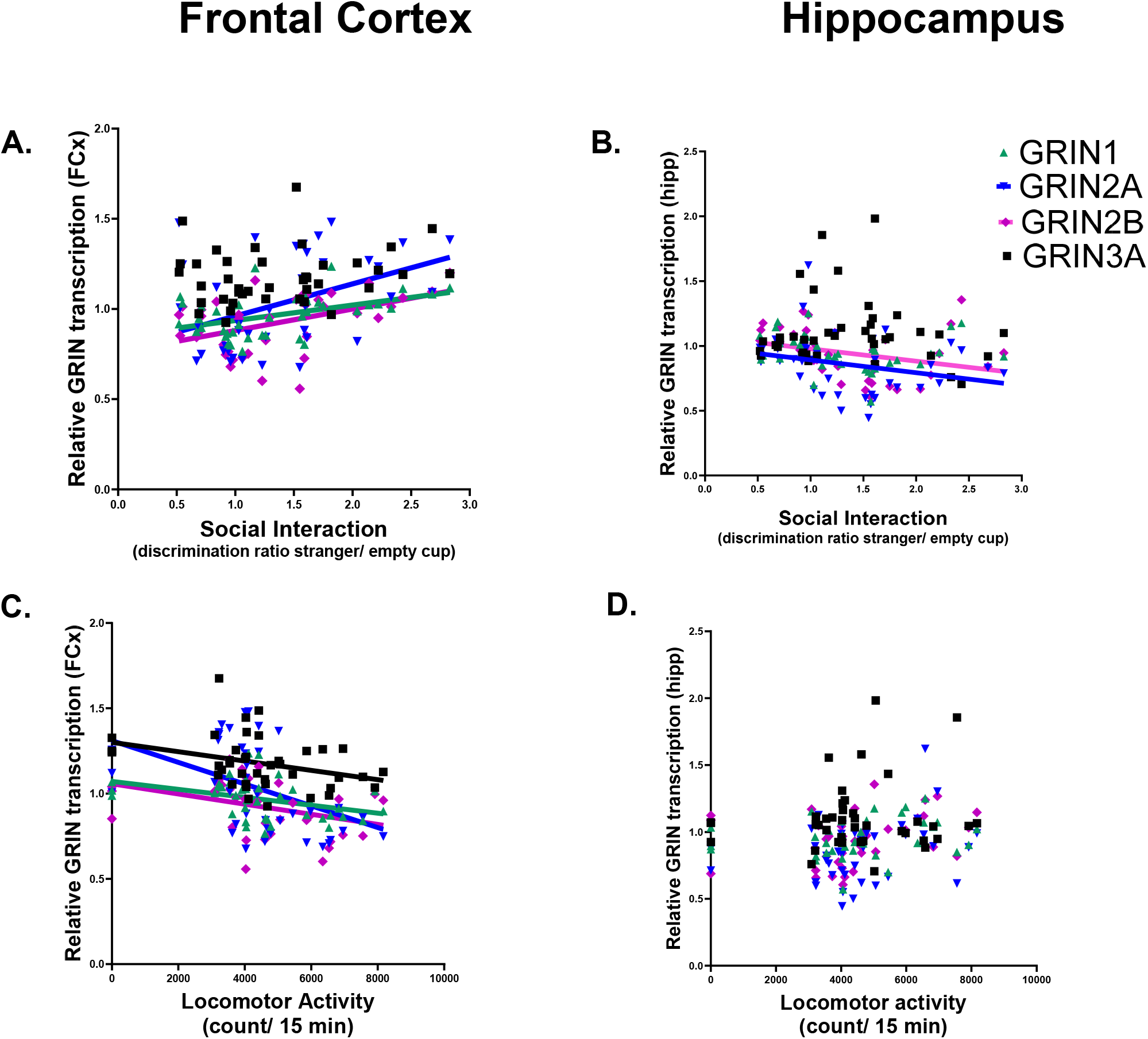
Social Interaction Behavior and Locomotor Activity correlate with GRIN transcription in the Hippocampus and Frontal Cortex. **(A)** Frontal Cortex GRIN transcription positively correlates with social interaction behavior. A Pearson’s test found a positive correlation between social interaction behavior and frontal cortex transcription of GRIN1 (r=0.45, p=0.01), GRIN2A (r=0.42, p=0.01), and GRIN2B (r=0.45, p=0.01). **(B)** Hippocampal GRIN transcription negatively correlates with social interaction behavior. Pearson’s test found a negative correlation between social interaction behavior and hippocampal transcription of GRIN2A (r=-0.33, p=0.05), and GRIN2B (r=-0.47, p=0.01). **(C)** Frontal Cortex GRIN transcription negatively correlates with locomotor behavior. Pearson’s test found a negative correlation between locomotor behavior and frontal cortex transcription of GRIN1 (r=-0.38, p=0.05), GRIN2A (r=-0.46, p=0.01), GRIN2B (r=-0.35, p=0.05), and GRIN3(r=- 0.34, p=0.05). **(D)** Hippocampal GRIN transcription does not correlate with locomotor behavior. A Pearson’s test found no significant correlation between Hippocampal GRIN transcription and locomotor activity. *Abbreviations:* Frontal Cortex, FCx; Glutamate Receptor Ionotropic NMDA, GRIN; Hippocampus, Hipp;

### Relationship between prenatal stress, GRIN transcription and locomotor activity

Levene’s test revealed homogeneity of variances in horizontal and vertical locomotor activity (p>0.05). Neither PRS nor drug treatment altered locomotor activity. Locomotor activity was not altered by PRS (F_1,34_=1.63, ns) or drug treatment (F_2, 34_=2.67, ns). There was no interaction between PRS exposure and drug treatment on locomotor activity (F_2, 34_=0.06, ns). The doses of clozapine and haloperidol produced marked sedation (clozapine) and catalepsy (haloperidol) one hour after injection in all mice, but these effects were temporary and not observed after 18 hours’ washout and therefore not present when we tested behavior.

We also found a negative correlation between horizontal locomotor activity and the transcription levels of GRIN genes in the frontal cortex (**Figure 2C**). There was no relationship between GRIN transcription in the hippocampus and locomotor activity (**Figure 2D**).

## Discussion

The data presented reveal that prenatal stress (PRS) significantly increases the transcription of NMDAR subunits in the hippocampus but not the frontal cortex. We observe a negative correlation between social behavior and NMDAR subunit transcription in the hippocampus, and in contrast, a positive correlation between social behavior and NMDAR subunit transcription in the frontal cortex. These data suggest that the circuitry expressing the NMDA receptor differs in response to stress in these brain regions.

We have previously reported that prenatal stress induces deficits of social interaction in adult mice (**Figure 1B**). Treatment with the atypical antipsychotic drug clozapine but not the first generation antipsychotic drug, haloperidol appeared to prevent these social deficits in the prenatally stressed mice (Bristow et al., 2021).

Antipsychotic treatment also significantly altered the transcription of NMDAR subunits, especially in the frontal cortex. This could be correcting the reduced glutamatergic responsiveness and overall circuit inhibition of the frontal cortex after prenatal stress (Fumagalli et al., 2009; Marchisella et al., 2021).

A previous study reported reduced expression of NR1 (GRIN1) and NR2B (GRIN2B) subunits in the hippocampal post-synaptic density of animals exposed to prenatal stress, resulting in lower numbers of functional NMDARs (Son et al., 2006). While our data appear to be inconsistent with these findings, this may be due to methodological differences between the two studies. Firstly, we measured mRNA abundance/ transcription while Son and colleagues measured protein abundance. There are advantages and disadvantages of each approach for example, measurement of RNA has greater molecular specificity than affinity-based assays using antibodies to measure protein abundance. Moreover, if the effects of stress on gene transcription is due to epigenomic modifications as indicated by studies of prenatal stress-induced changes in DNA methylation (Dong et al., 2016a), then transcription of mRNA is the next, more immediate step in the gene expression process, and is followed by translation to protein. A second methodological difference was in the strains of mice tested in the studies. We have tested Swiss Webster mice while Son et al. tested ICR mice (Son et al., 2006). Although these strains have common founders, Swiss Webster mice have a disrupted in schizophrenia-1 (DISC1) mutation while ICR mice are homozygous for the wild-type DISC1 gene. A recent meta-analysis reported that mutations in the human DISC1 gene are strongly associated with schizophrenia (Wang et al., 2018). DISC1 mutations modify NMDAR dynamics, including increasing the density of NMDAR synaptic vesicles and the expression of NMDARs at the synaptic surface. DISC1 regulates the exit of GluN1 (encoded by GRIN1) from the endoplasmic reticulum. In addition, DISC1 alters the transport of GluN2A (encoded by GRIN2A) and GluN2B (encoded by GRIN2B) (Harrison and Weinberger, 2005; Carter, 2006; Wu et al., 2013). Overall, it is possible that the DISC1 mutation in the Swiss Webster strain might increase the susceptibility of this mouse strain to the pathology of prenatal stress through modified regulation of NMDAR biogenesis. This could be a gene x environment dynamic that could recapitulate some aspects of the etiology of stress-induced pathology in the hippocampus that occurs in schizophrenia and other disorders associated with social deficits. Son et al. also used longer stress exposure conditions in their mice, which may have been necessary with the ICR mouse strain but not in the genetically vulnerable Swiss Webster strain. In contrast, studies of stress exposure in adult rats reveal increased transcription of NMDA receptor subunits in the hippocampus, including GluN2A and GluN2B (Calabrese et al., 2012). This stress-induced increase in NMDA receptor subunits is thought to contribute to enhanced excitability and vulnerability of hippocampal neurons (Schwendt and Jezova, 2000). These findings collectively suggest that stress can lead to a dysregulation of glutamate receptor subunits in the hippocampus, potentially contributing to the pathophysiology of stress-related disorders.

### Limitations of the Study

Our study has limitations that are common in neurobiological analyses. We have focused the experimental plan on males, because social deficits associated with prenatal stress are observed in males and not females (Weinstock, 2008). Nevertheless, the underlying mechanisms of resilience to prenatal stress in females are of importance and could provide insights into neuroprotective mechanisms. We focused analyses on the frontal cortex and hippocampus, and future studies should include investigations of subregions within the frontal cortex and hippocampus, in addition to the amygdala, to determine the regional specificity of the underlying pathological mechanisms triggered by prenatal stress.

*In summary*, this study illustrates the importance of the impact of prenatal stress on the glutamate system in the hippocampus. The data indicate that prenatal stress alters the transcription of NMDAR subunits in the hippocampus and not frontal cortex. Altered transcription of NMDAR subunits is likely to have a profound impact on excitatory transmission, and cognitive behaviors associated with the corticolimbic circuit. This impact on the corticolimbic circuit could be the novel molecular pathway through which prenatal stress leads to the long-term deficits in social behavior that are common to several psychiatric disorders. More detailed analyses of molecular pathways associated with glutamatergic transmission in the hippocampus could provide novel targets for the development of treatments for stress-associated psychiatric disorders that include symptoms of social withdrawal.

## Acknowledgements

Funded by the Department of Molecular Pharmacology and Neuroscience at Loyola University Chicago.

